# Transcriptomic response of *Caenorhabditis elegans* expressing human Aβ_42_ gene treated with Salvianolic acid A

**DOI:** 10.1101/2020.05.28.120485

**Authors:** Chee Wah Yuen, Mardani Abdul Halim, Nazalan Najimudin, Ghows Azzam

## Abstract

Alzheimer’s disease (AD) is a brain disease attributed to the accumulation of extracellular senile plaques comprising β-amyloid peptide (Aβ). In this study, a global transcriptomic analysis of the response of transgenic *Caenorhabditis elegans* worms expressing full length human Aβ_42_ gene towards Salvianolic acid A (Sal A) was analysed. Antioxidant response genes, namely *gst-4*, *gst-10*, *spr-1* and *trxr-2*, were upregulated. The production of Aβ_42_ caused oxidative stress and the antioxidant response genes possibly provide defence to the strain. The gene product of *trxr-2* also functionally interacts with the defence system and has a role in life span. Genes involved in replication, reproduction, immune response to microbes and antimicrobial activities were also upregulated. Exposure to Sal A also increased the rate of reproduction of nematodes, and heightened its immunological protection system towards microorganisms. In contrast, genes responsible for locomotion, ligand-gated cation channel, embryonic and postembryonic development, and neuromodulation of chemosensory neurons were significantly down-regulated. As an effector, Sal A might conceivably reduce the movement of the worm by interfering with neuronal transmission and embryonic and post-embryonic development.

## Introduction

Alzheimer’s disease (AD) is a chronic, progressive brain disorder which affects middle or elderly population. The disease eventually will lead to the loss of thinking ability and slowly getting worse over time until it become severe enough to interfere with daily task. AD is the most common type. The number of demented patients was predicted to double every 20 years from 24.3 million in 2001 to 42.3 million in 2020 to a whooping number of 81.1 million in 2040 worldwide [1].

AD can be explained using the amyloid cascade hypothesis. The proposed amyloid hypothesis was supported with several evidences. The first evidence was the discovery of senile plaques (SPs) and neurofibrillary tangles (NFTs) by a physician, Dr Aloi Alzheimer, when autopsy was performed on AD patient’s brain in 1907 (reviewed by Teplow *et al*., 2012) [2]. This was followed by significant findings of amyloid-beta (Aβ) within SPs [3], discovery of amyloid precursor protein (APP) gene sequence [4] mutations [5] which led to the proposed amyloid cascade hypothesis [6]. Briefly, the amyloid cascade hypothesis encompasses the scission of APP by β-secretase and γ-secretase to form Aβ peptides, followed by aggregation of Aβ oligomers to produce SPs and eventually causing toxicity to the brain cells by oxidative stress.

Based on the amyloid cascade hypothesis, a search for potential therapeutics that inhibits the Aβ production had been initiated. Among the potential therapeutics that hinders Aβ production are drugs that contain anti-Aβ aggregation properties to disrupt the formation of SPs and antioxidant effects to decrease the oxidative stress caused by Aβ. Since there is no disease modifying drugs for AD, the current drugs that are available can only delay the onset of AD [7]. However, there are drawbacks for these drugs in the effort to modify the AD pathogenesis and the deleterious side effects of the present drugs, thus, there is a need to search for new potential disease modifying AD drugs.

Salvianolic acid A (Sal A) is one of the major water-soluble compounds in the water Danshen extract besides salvianolic acid B and danshensu [8]. Sal A have exhibited its antioxidant properties by inhibiting ROS production due to H_2_O_2_ induction, and significantly scavenging HO. that was produced in phorbol myristate acetate-stimulated rat neutrophils [9, 10]. Studies had demonstrated that Sal A has the ability to improve memory impairment when the compound was injected intravenously in mice [11]. It was also reported that 10 mg/kg Sal A that was intravenously injected into the mice can inhibit cerebral lipid peroxidation and clear free HO. radicals. Hence, it can be deduced that there is a relationship between its antioxidant properties with its improving effects on memory impairment induced by cerebral ischemia-reperfusion in mice [11].

Our previous study using *C. elegans* expressing human Aβ42 gene have shown that Sal A has the ability to defer paralysis of the worm. Besides, Sal A treatment to the same worm also inhibits the development of Aβ fibrils plus it decreased the Aβ-induced ROS [12]. In this study, transgenic *Caenorhabditis elegans* carrying Aβ_42_ gene was used as model organism to study the global effect of Sal A towards levels of *C. elegans* transcripts via RNA-seq analysis.

## Methods and Materials

### C. elegans strains and maintenance

Transgenic *C. elegans* strain GMC101(dvIs100[unc-54p::A-beta-1-42::unc-54 3’-UTR + mtl-2p::GFP) and *E. coli* OP50 strain was kindly provided by the Caenorhabditis Genetics Center (CGC), University of Minnesota (Minneapolis, MN, USA). The transgenic C. elegans were maintained at 16 °C on nematode growth medium (NGM) seeded with *E. coli* OP50 bacteria. GMC101 is a transgenic strain that expressed the human Aβ_42_ gene.

### RNA extraction, quality and quantity evaluation

*C. elegans* strain GMC101 worms cultured on NGM plates supplemented with 100μg/ml Sal A was used for transcriptomic study. For each treatment, approximately 5000 adult worms were harvested after 32 hours of upshift from 16 °C to 25°C to induce Aβ expression intended to cause paralysis to occur. They were harvested by washing with M9 medium to remove *E. coli* OP50 cells. RNA isolation was conducted as described by He (2011) [13] using AccuZol (Bioneer, Korea). The RNA samples were treated with DNase (Macherey-Nagel, Germany) to remove any DNA present in the samples and the RNA samples were repurified using NucleoSpin RNA Clean-up XS (Macherey-Nagel, Germany). The total RNA was quantified using agarose gel electrophoresis and NanoPhotometer® spectrophotometer (IMPLEN, CA, USA). RNA concentration was measured using Qubit® RNA Assay Kit in Qubit® 2.0 Flurometer (Life Technologies, CA, USA) and RNA integrity was assessed using the RNA Nano 6000 Assay Kit of the Bioanalyzer 2100 system (Agilent Technologies, CA, USA). The same protocol was also used to prepare RNA samples for quantitative real-time PCR (qPCR).

### RNA-seq library preparation

For the library construction, standard Illumina protocol was employed involving fragmentation of mRNA, synthesis of double stranded cDNA, polyadenylation, adapter ligation and library size selection (150-200 bp). The libraries were generated using NEBNext® Ultra™ RNA Library Prep Kit for Illumina® (NEB, USA). Briefly, mRNA was purified from total RNA using poly-T oligo-attached magnetic beads. . Fragmentation was carried out using divalent cations under elevated temperature in NEBNext First Strand Synthesis Reaction Buffer. First strand cDNA was synthesized using random hexamer primer and M-MuLV Reverse Transcriptase. The second strand cDNA synthesis was performed using DNA Polymerase I. The remaining overhangs were converted into blunt ends via exonuclease/polymerase activities. After adenylation of 3’ ends of DNA fragments, NEBNext Adaptor with hairpin loop structure were ligated to prepare for hybridization. In order to select cDNA fragments of 150~200 bp in length, the library fragments were purified with AMPure XP system (Beckman Coulter, Beverly, USA) and adapter was ligated. Final libraries were generated by PCR and the library was assessed by Agilent Bioanalyzer 2100 system and subsequently used for RNA-seq. Raw data generated was cleaned and trimmed by removing the adapter as well as removing the low quality reads.

### Mapping of reads to the reference genome

The reference genome and gene model annotation files were downloaded from the genome websites [ftp://ftp.ensembl.org/pub/release-76/fasta/caenorhabditis_elegans/dna] and [ftp://ftp.ensembl.org/pub/release-76/gtf/caenorhabditis_elegans], respectively. An index of the reference genome was built using Bowtie v2.2.8.0 [14, 15] and paired-end clean reads were aligned to the reference genome using TopHat v2.1.1 [16]. The programme HTSeq v0.6.1 was used to count the reads numbers mapped to each gene [17]. The FPKM value of each gene was calculated based on the length of the gene and reads count mapped to this gene. FPKM is the expected number of Fragments Per Kilobase of transcript sequence per Millions base pairs sequenced. It considers the effect of sequencing depth and gene length for the reads count at the same time, and is currently the most commonly used method for estimating gene expression levels [18].

### Differential expression analysis

The DESeq R package version 1.18.0 was used to perform the differential expression analysis of the two experimental conditions (with two biological replicates per condition) [19]. The read counts of each sequenced library were adjusted by the edgeR program package through one scaling normalized factor [20]. Differential expression analysis of the two conditions was performed using the DEGSeq R package (1.20.0) [21]. The P values were adjusted using the Benjamini & Hochberg method. Corrected P-value of 0.005 and log2 (Fold change) of 1 were set as the threshold for significantly differential expression [22].

### GO and KEGG enrichment analysis of differentially expressed genes

Gene Ontology (GO), enrichment analysis of differentially expressed genes was implemented by the GOseq R package, in which gene length bias was corrected. GO terms with corrected Pvalue less than 0.05 were considered significantly enriched by differential expressed genes [23]. KOBAS software was used to test the statistical enrichment of differential expression genes in KEGG pathways [24].

### Real-time PCR validation of transcriptomic results

To validate the bioinformatics analysis of differentially express patterns, 10 genes (five up- and five downregulated genes) were chosen. The primers of the selected genes were designed using Primer 3 software (https://www.ncbi.nlm.nih.gov/tools/primer-blast/) and synthesized commercially (Integrated DNA Technologies; Singapore) (supplementary 1). Normalization was carried out against two reference genes which were *tba-1* (tubulin, alpha family member gene), and *cdc-42* (cell division cycle related gene) [25]. All reactions were conducted with 3 biological and technical replicates.

## Results

### Clean reads mapping

A total of 48540212, 47260368 and 48116402 raw reads were generated from treated samples for replicate 1 (Treat_S1), replicate 2 (Treat_S2) and replicate 3 (Treat_S3), respectively. For untreated samples (control), a total of 42927272, 40774690 and 40110332 of raw reads were generated for replicate 1 (CT1), replicate 2 (CT2) and replicate 3 (CT3), respectively. After QC, clean reads obtained were 46931624, 45680280 and 46326584 for treated samples while 41562272, 38370118 and 38869146 reads were obtained from control samples.

The cleaned reads were mapped to *C. elegans* genome. For control samples CT1, 96.7% of the reads were mapped to exon region, 3.2% intergenic and 0.1% intron. For CT2, 92.4% were mapped to exon, 7.5% intergenic and 0.1% intron. For CT3, 95.4% were mapped to exon, 4.5% intergenic while 0.1% to intron region. On the other hand, for treated samples Treat_S1, 85.3% of the cleaned reads were mapped into exon region, 14.6% intergenic and 0.1% intron. For Treat_S2, 91.6% mapped to exon, 8.3% mapped to intergenic and 0.1% intron. Finally, for Treat_S3, 91.2% of the cleaned reads were mapped to exon region, 8.7% mapped to intergenic and 0.1% to intron. Fig. 1 and Fig. 2 summarized the statistic of mapped cleaned reads to *C. elegans* genome. Pearson correlation was used to determine relation of treated and control samples (Fig. 3). From the graph, all samples in treated showed high similarity among replicates and the same correlation was also observed for control samples. After the reads were mapped into chromosome, they were internally normalized to minimize the effect of sequencing depth and gene length for the reads count at the same time. Fig. 4 showed the distribution of control samples vs treated samples in FPKM.

**Fig. 1.**
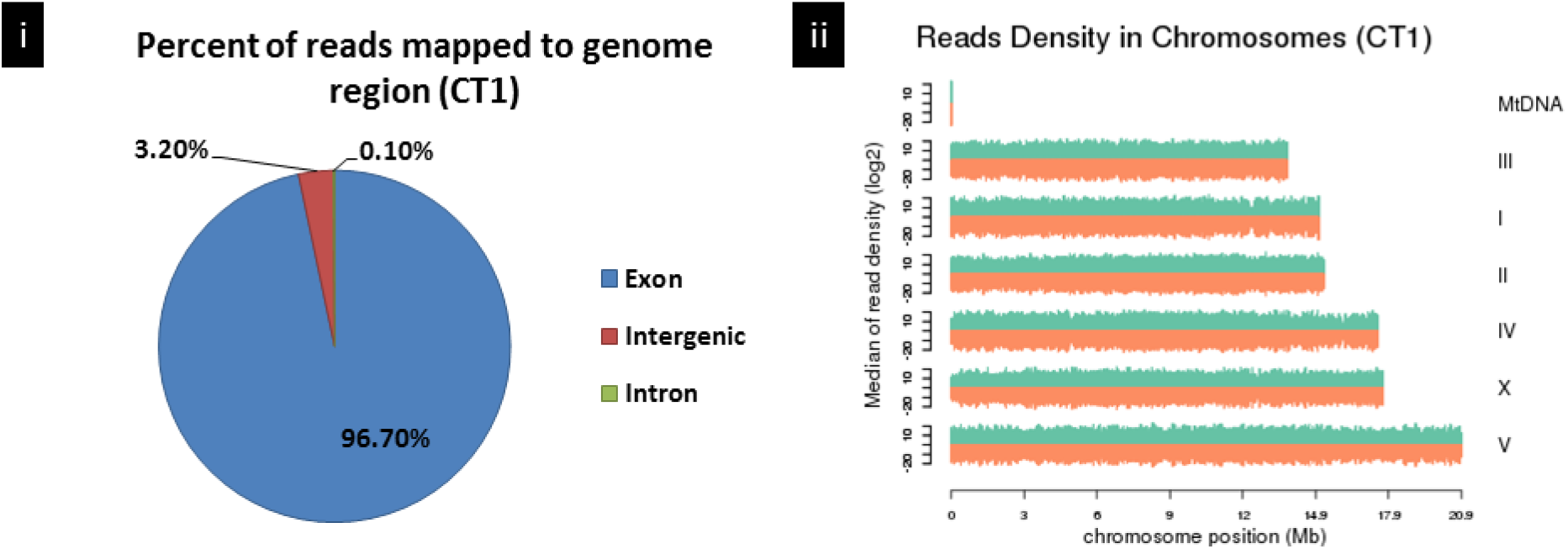
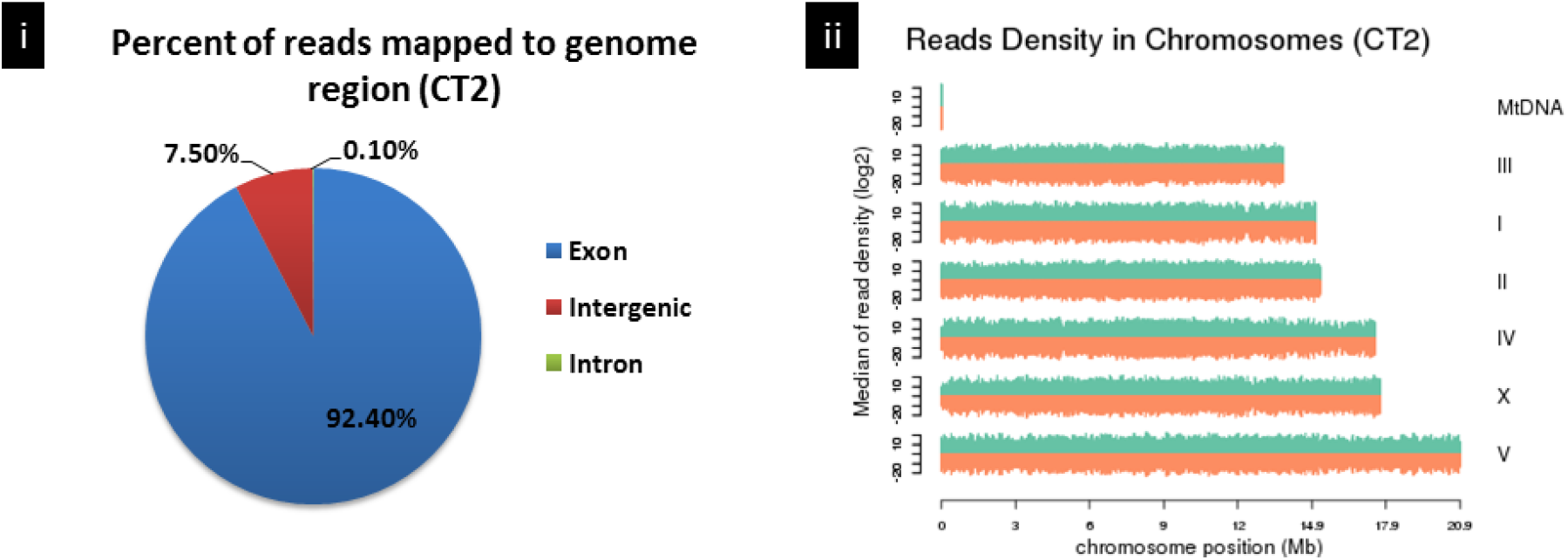
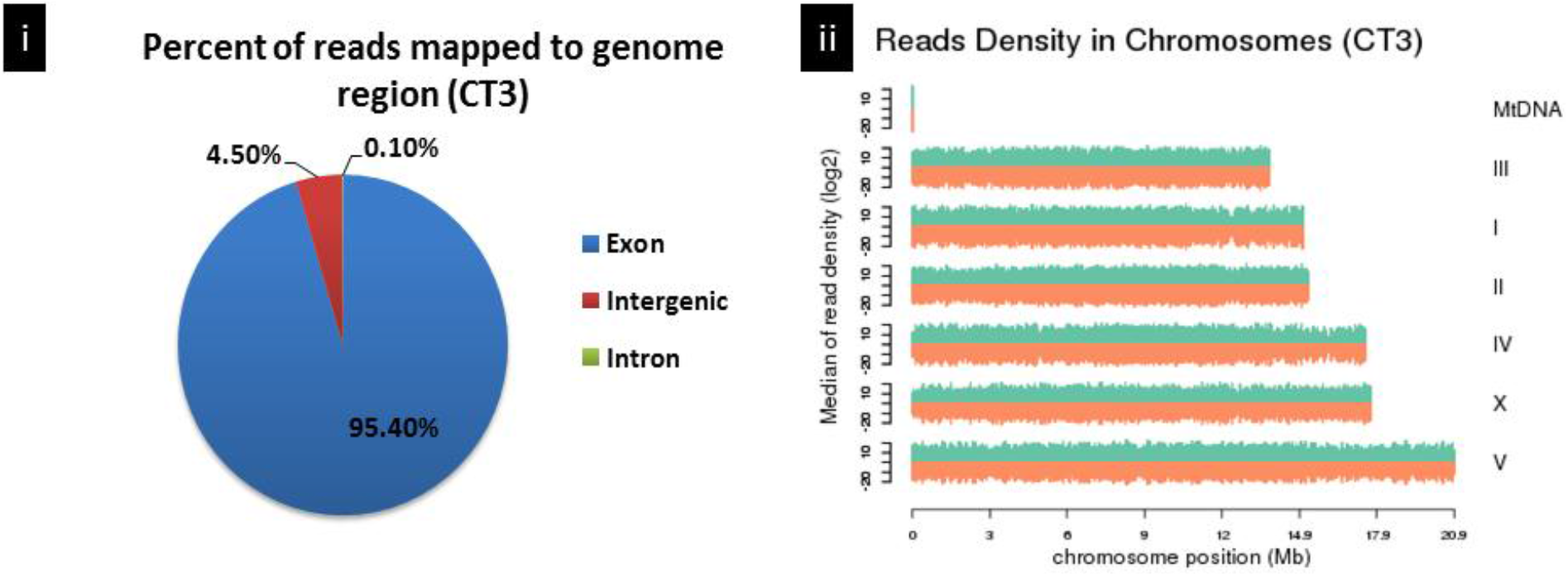
Statistic of mapped reads of CT samples to *C. elegans* chromosome. Fig. 1A, 1B and 1C represent the replicates of CT samples. The lowest mapped reads (neglecting MtDNA) was chromosome III followed by I, II, IV and X. The highest reads density was observed in chromosome V. Similar pattern was observed for all replicates.

**Fig. 2.**
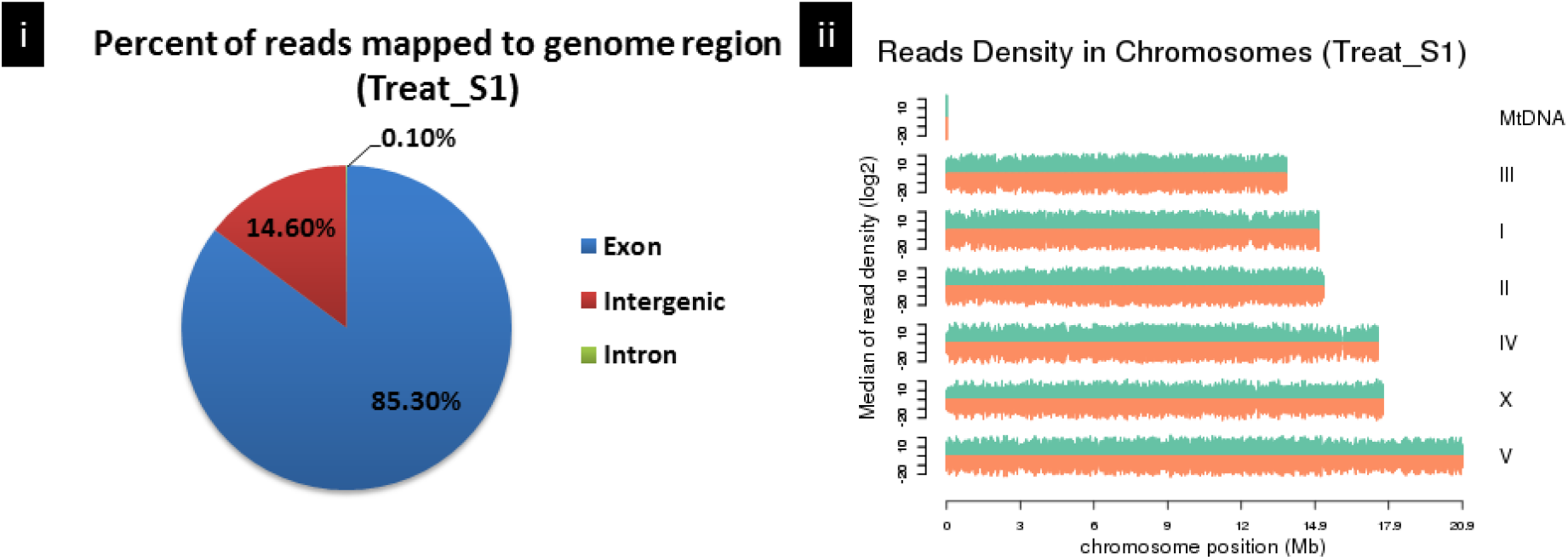
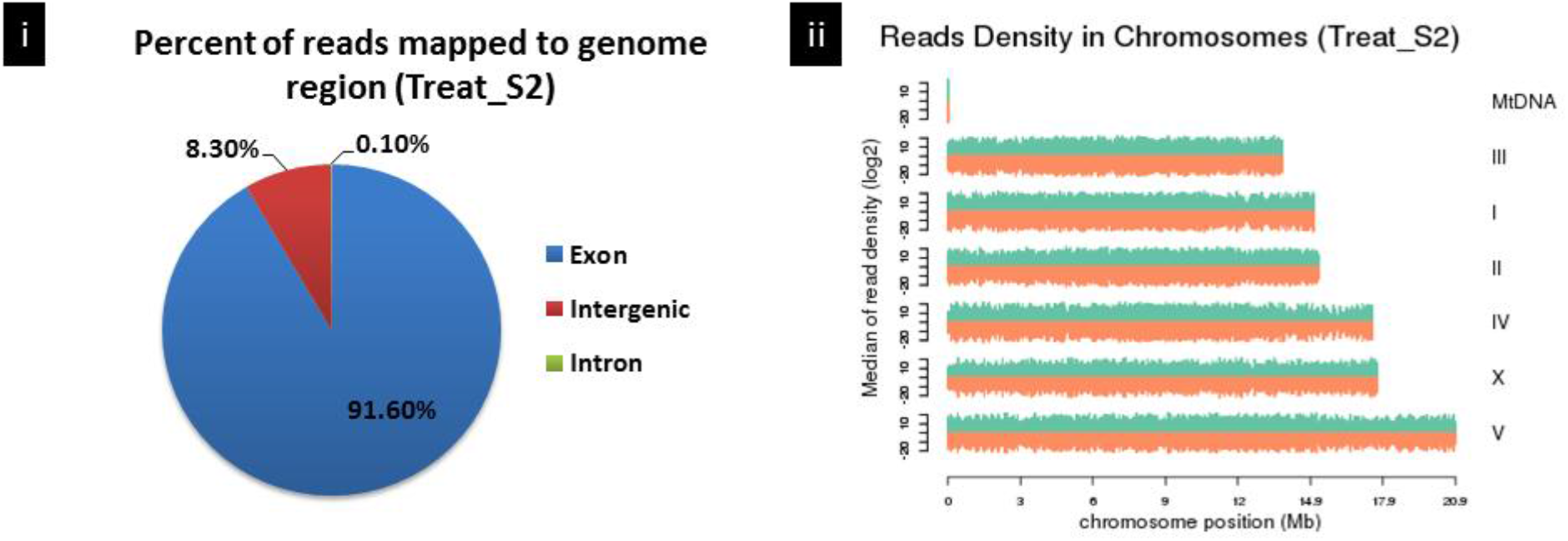
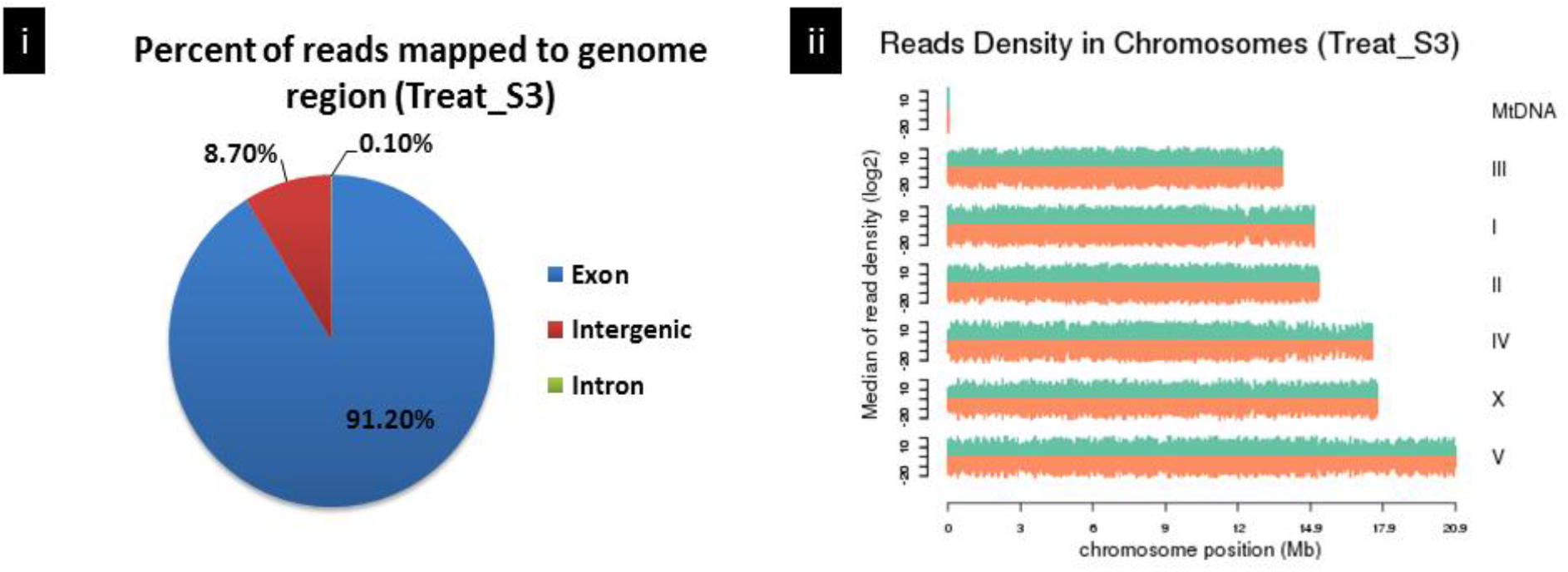
Statistic of mapped reads of treated samples to *C. elegans* chromosome. Fig. 2A, 2B and 2C represent the replicates of treated samples. The lowest mapped reads (neglecting MtDNA) was chromosome III followed by I, II, IV and X. The highest reads density was observed in chromosome V. Similar pattern was observed for all replicates.

**Fig. 3.**
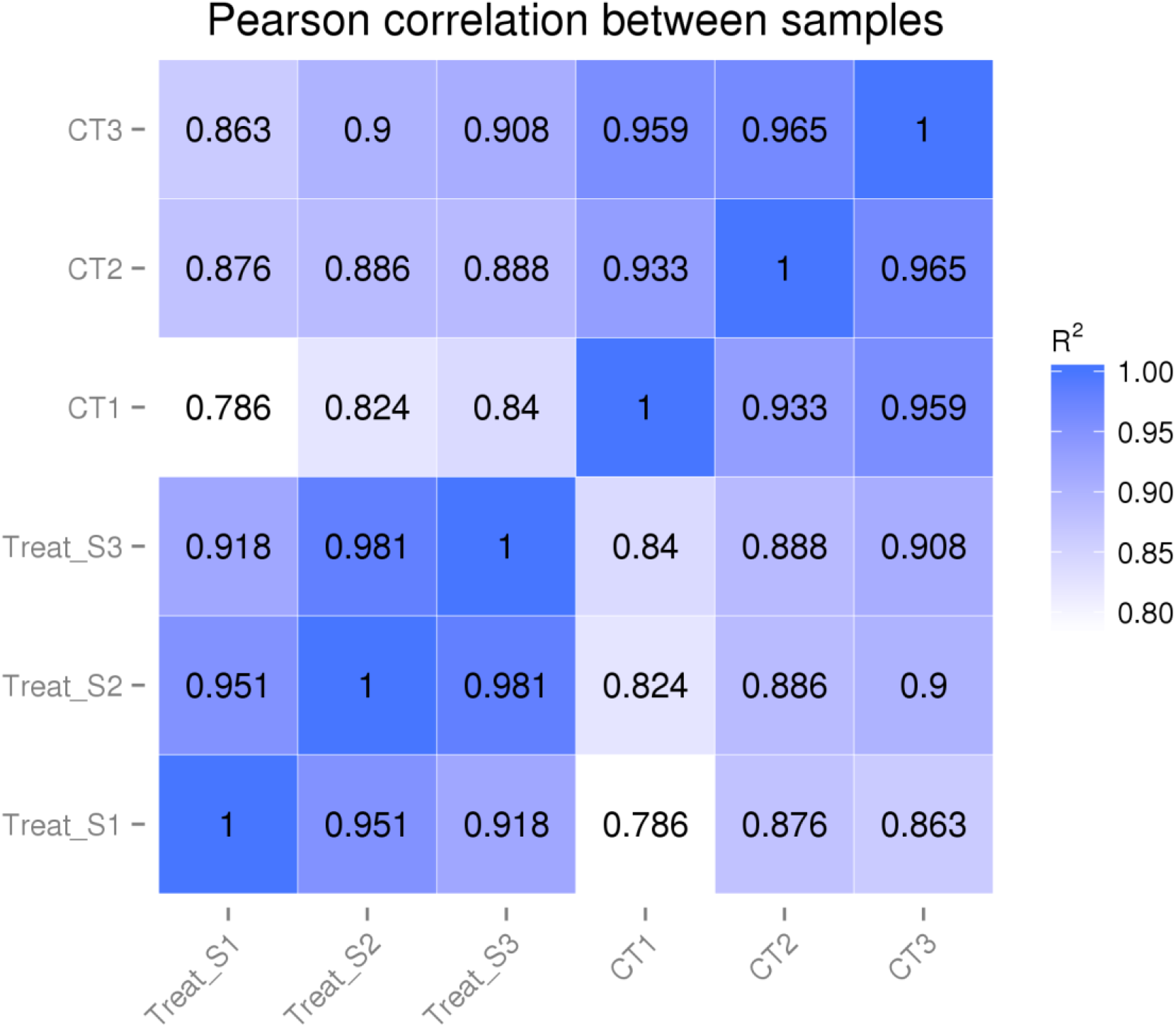
Pearson correlation graph of control vs treated samples.

**Fig. 4.**
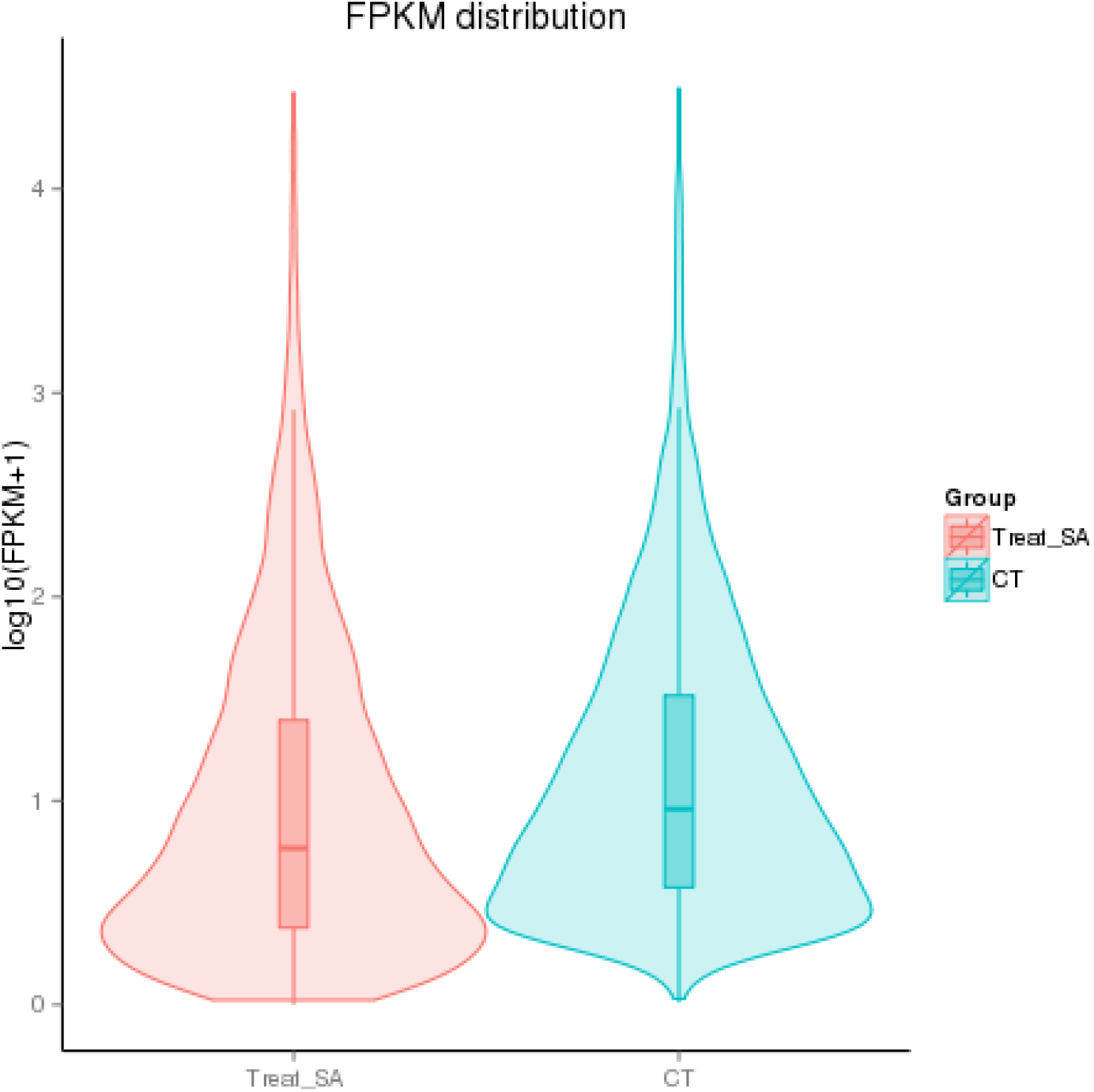
Distribution of FPKM value of control vs treated samples.

### Identification of differentially expressed genes

Based on the transcriptomic studies, Sal A-treated and untreated *C. elegans* GMC101 strain showed differences in global gene expression. The treated *C. elegans* showed down-regulation of 2020 genes (Supplementary 2) and up-regulation of 1435 genes (supplementary 3) when variations of at least 2-fold were considered. The differentially expressed genes were clustered together (control vs treated) and were presented in the form of heat map (Fig. 5).

**Fig. 5.**
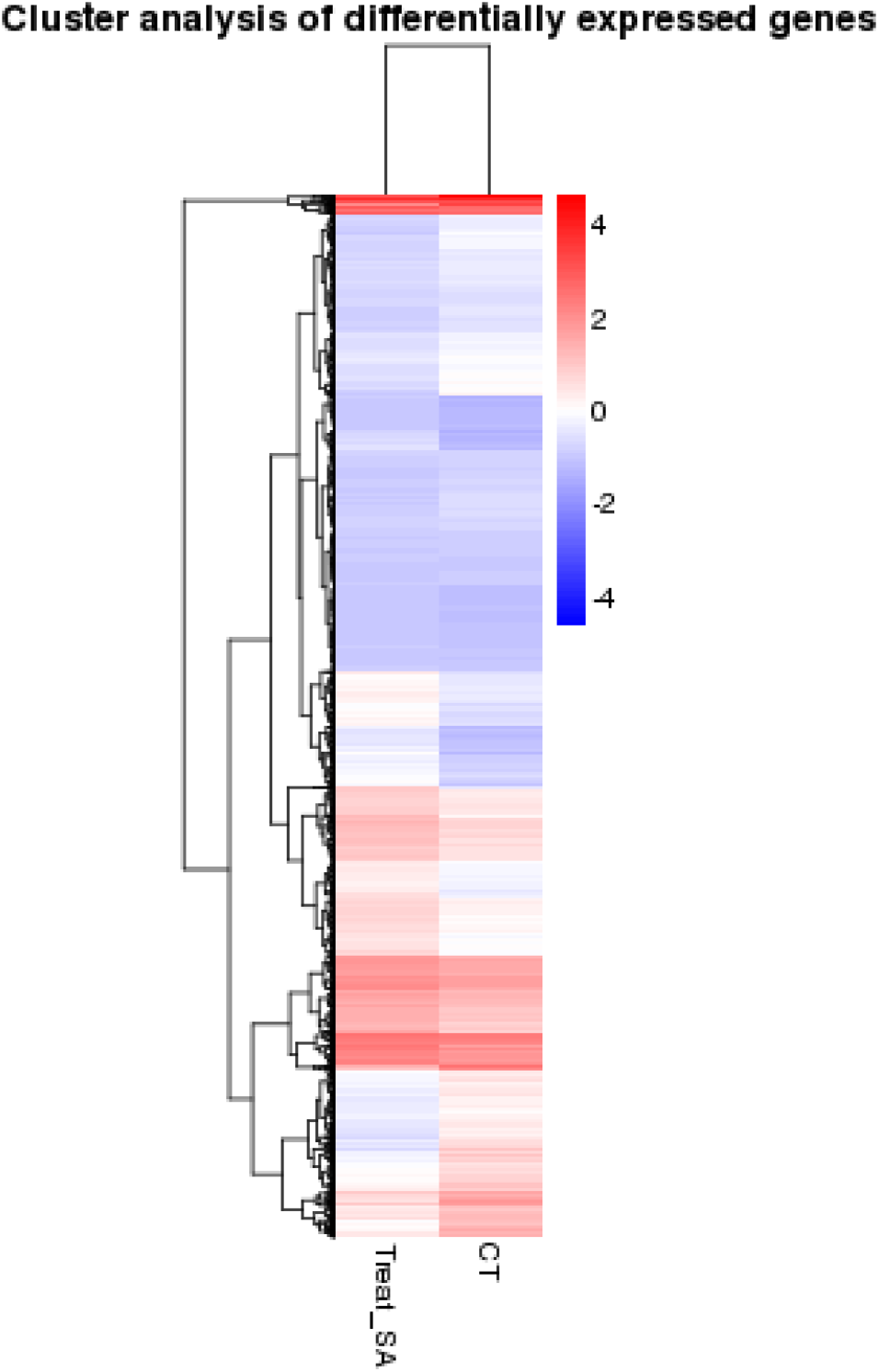
Cluster analysis of differentially expressed genes.

### Sal A affected homologues associated with Alzheimer’s disease in human

Based on the transcriptomic results, two homologues that are related to Alzheimer’s disease in human were affected by Sal A (Fig. 6). The gene *trxr-2*, a *putative thioredoxin reductase gene,* was upregulated while *ptl-1,* a homolog of the MAP2/MAP4/tau family in the nematode, was downregulated.

**Fig. 6:**
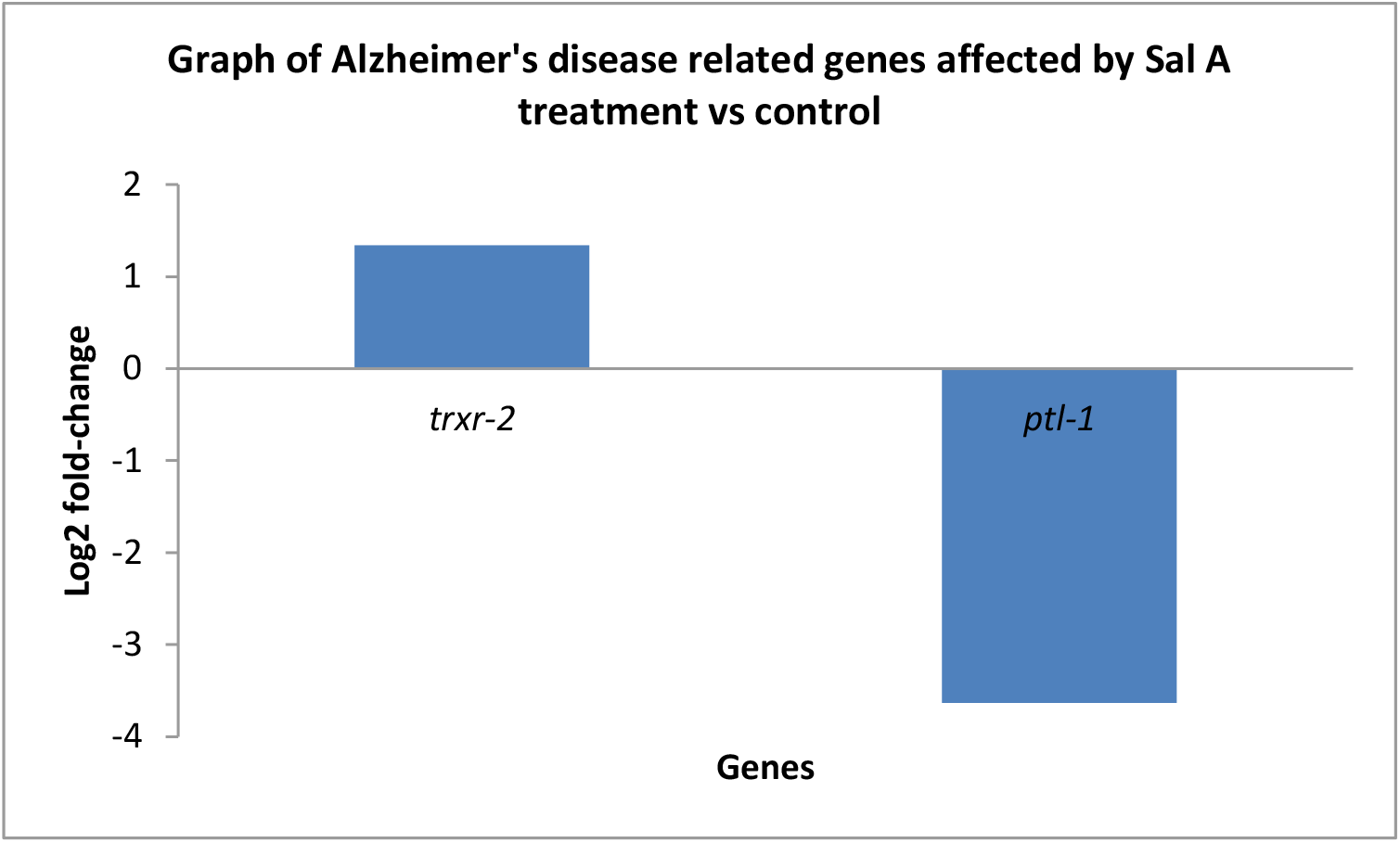
Genes that were related to Alzheimer’s disease were affected by Sal A.

### Sal A upregulated antioxidant response genes in the transgenic C. elegans GMC101 strain

Among the differentially expressed genes, three antioxidant response genes were identified, namely, *sod-1*, *gst-4*, *gst-10* and *spr-1*. These genes were affected when *C. elegans* strain GMC101 was fed with 100 μg/ml Sal A compared to the unfed nematodes. The gene expressions of antioxidant response genes such as *gst-4*, *gst-10* and *spr-1* were enhanced as can be seen in Fig. 7.

**Fig. 7:**
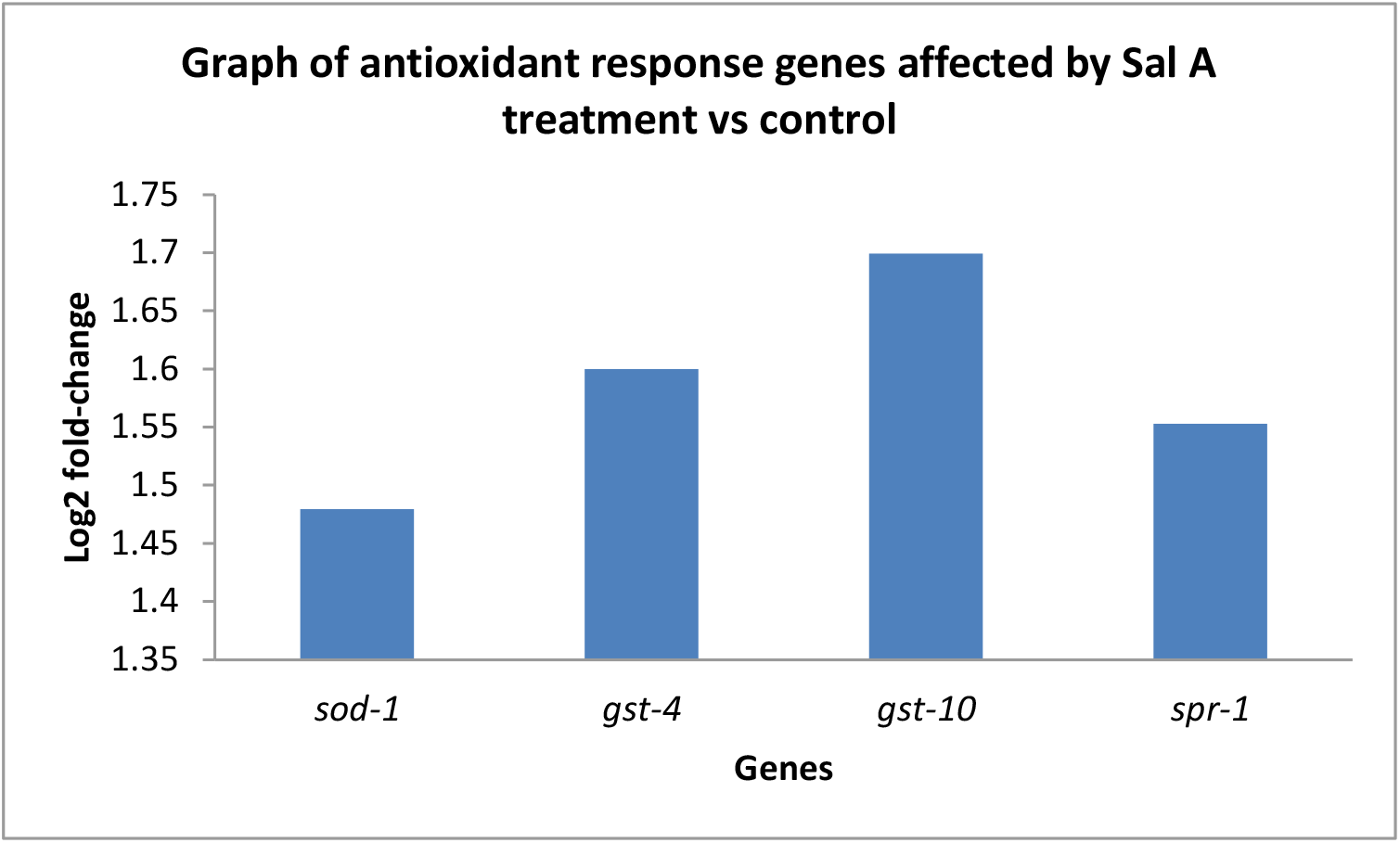
Antioxidant response genes that were affected by Sal A.

### GO enrichment analysis

All the 3455 differentially expressed genes (DEGs) were subsequently analyzed by using GO enrichment analysis. The genes were classified into three broad categories of biological process, cellular component and molecular function (Fig. 8).

**Fig. 8:**
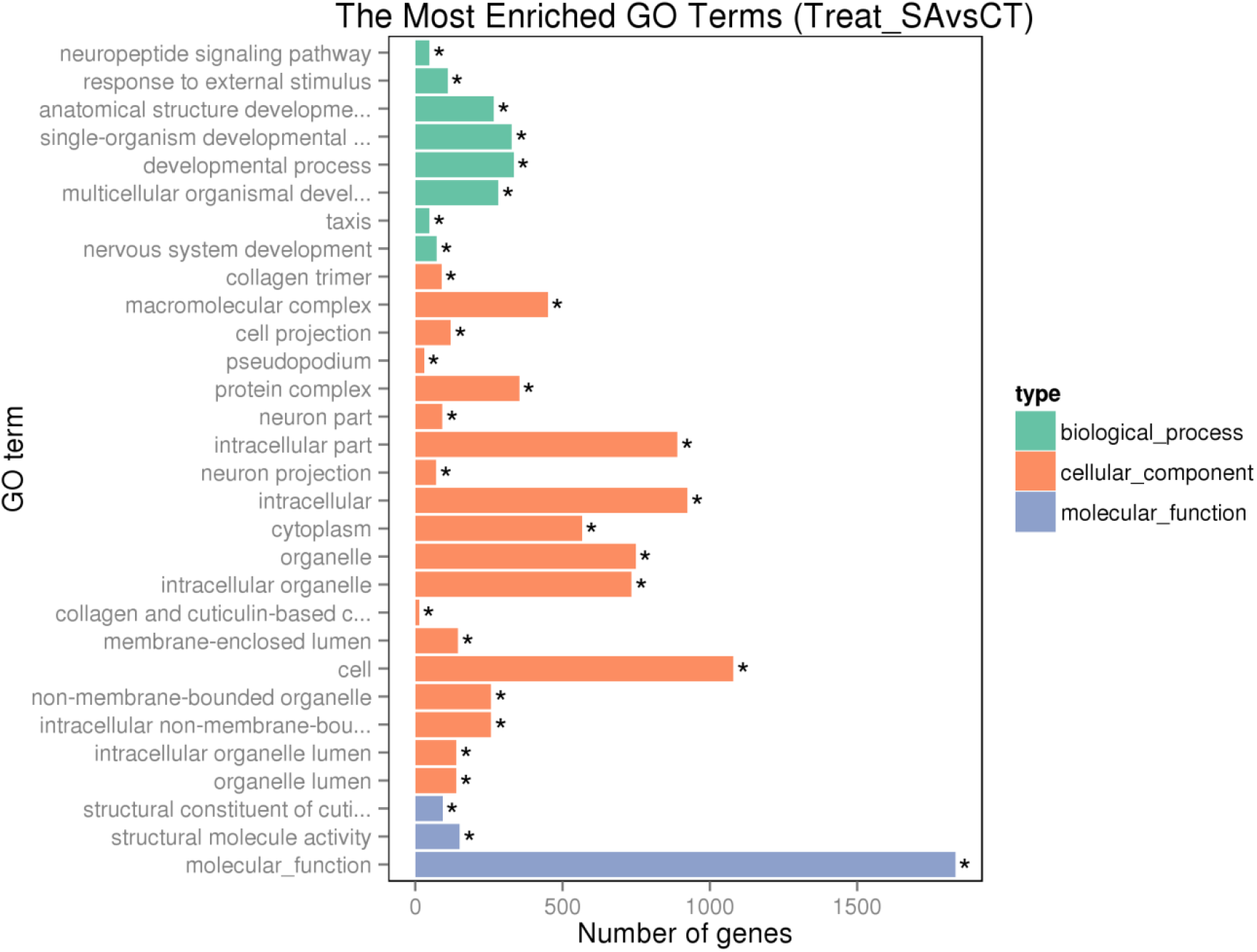
Enriched GO classification.

For down-regulated GO, the subcategories that classified under “biological process” were neuropeptide signalling pathway, response to external stimulus, nervous system development, cell communication, response to stimulus, single organism signalling, signalling, neuron differentiation, taxis, generation of neurons, locomotion, neurogenesis, multicellular organismal development, chemotaxis, anatomical structure development, collagen and cuticulin-based cuticle development and system development. As for “cellular component” category, the main subcategories for down-regulated GOs were collagen trimer, neuron part, extracellular region, neuron projection, cell projection, extracellular region part, synapse, somatodendritic compartment and axon. The major subcategories of the “molecular function” category for down-regulated GOs induced by Sal A include structural constituent of collagen and cuticulin-based cuticle, structural molecule activity, receptor binding and hormone activity. Fig. 9 summarized the most enriched down-regulated GO terms. The total down-regulated GO analysis was presented in Supplementary 4.

**Fig. 9:**
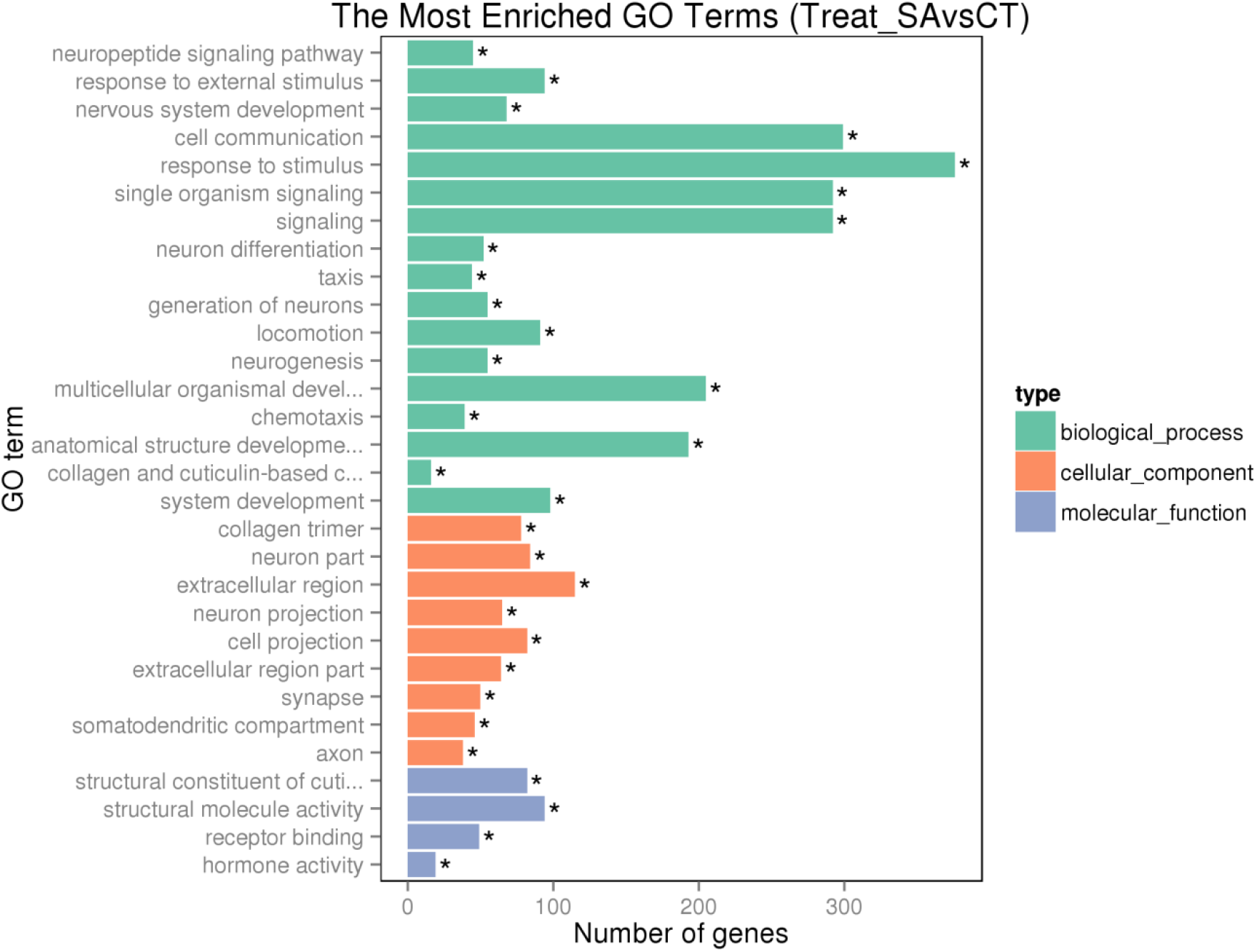
The most enriched down-regulated GO.

The most enriched GO categories that were classified as up-regulated were “biological process” and “cellular component”. For “biological process”, the subcategories involved were cellular nitrogen compound, cellular metabolic process, nitrogen compound metabolic process, metabolic process, organic substance metabolic process, RNA processing and primary metabolic process. For “cellular component”, the subcategories involved were intracellular part, intracellular, intracellular organelle, organelle, cytoplasm, intracellular organelle part, cell part, intracellular membrane-bounded organelle, cell, membrane-bounded organelle, organelle part, macromolecular complex, cytoplasmic part, non-membrane-bounded organelle, intracellular non-membrane-bounded organelle, membrane-enclosed lumen, intracellular organelle lumen, organelle lumen, nuclear part, nuclear lumen, intracellular ribonucleoprotein complex, ribonucleoprotein complex and mitochondrion. Fig. 10 summarized the most enriched up-regulated GO terms. The total up-regulated GO analysis was presented in Supplementary 5.

**Fig. 10:**
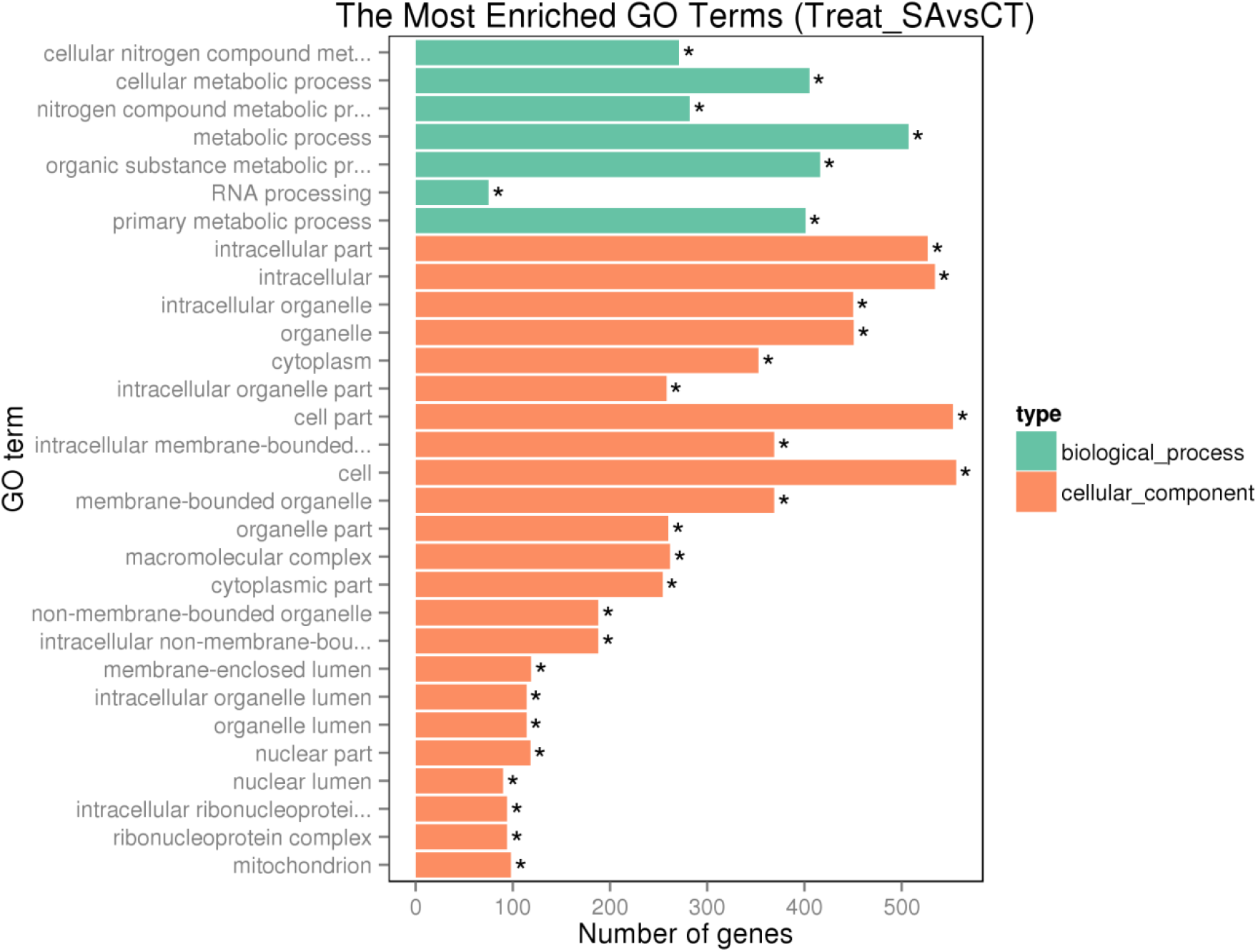
The most enriched up-regulated GO.

### KEGG pathway enrichment analysis

The assembled DEGs were also subsequently searched against the Kyoto Encyclopedia of Genes and Genomes (KEGG) database to determine which genes are involved in metabolic pathways. Fig 11 showed the statistic of overall KEGG pathway enrichment. The 4 significant up-regulated pathways (p<0.05) induced by Sal A were those involving RNA polymerase (14 genes), pyrimidine metabolism (23 genes), ribosome (35 genes) and oxidative phosphorylation (29 genes). The overall statistic of KEGG pathway enrichment for up-regulated pathway was presented in Fig. 12. The significantly down-regulated pathways (p<0.05) included those involving Wnt signalling (17 genes) and TGF-beta signalling (10 genes). Fig. 13 summarized the statistic of down-regulated pathway enrichment in KEGG. Table 1 showed the hyperlink to the KEGG pathway. The total KEGG analysis was presented in Supplementary 6 for up-regulated and Supplementary 7 for down-regulated pathways.

**Table 1.**
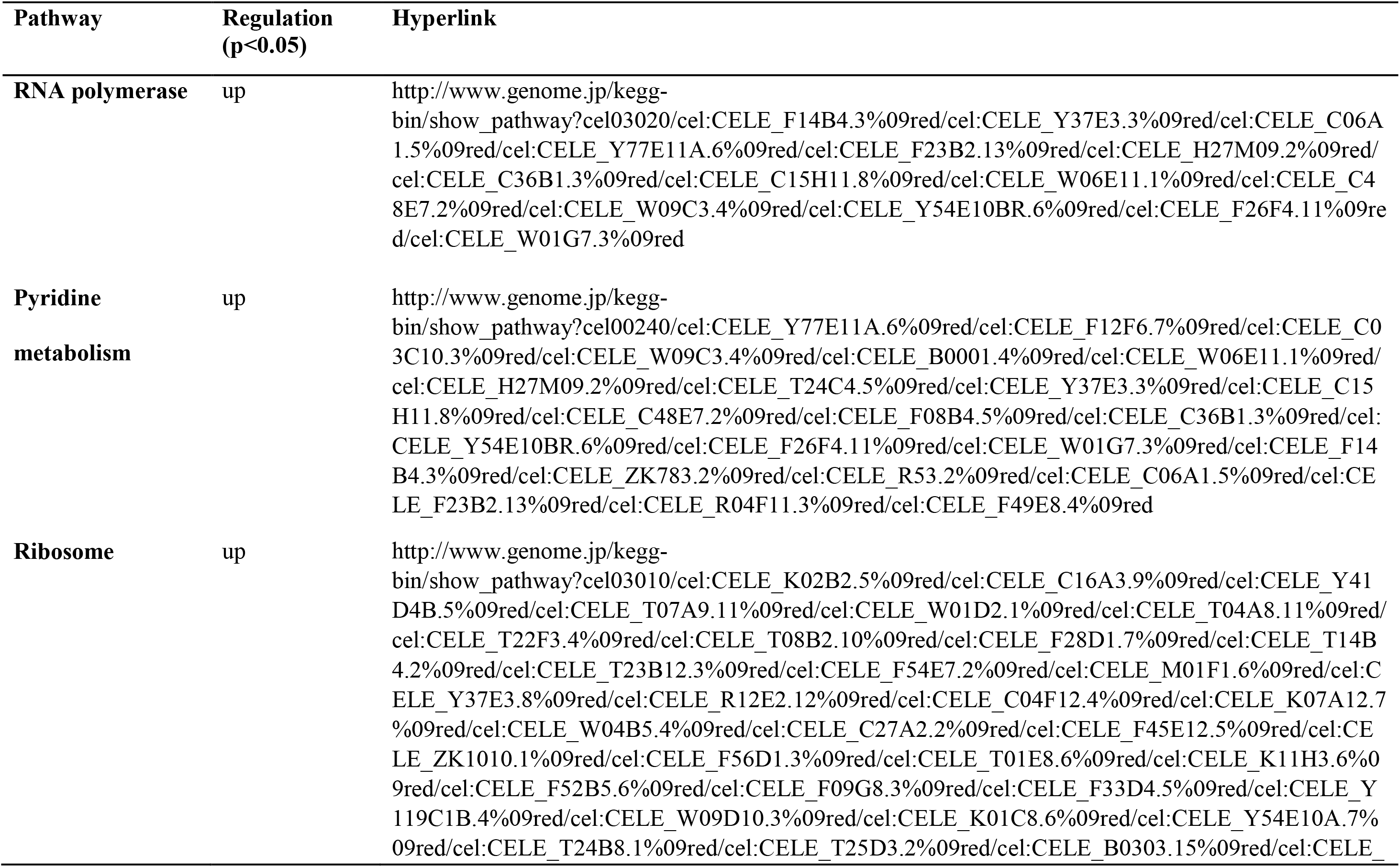
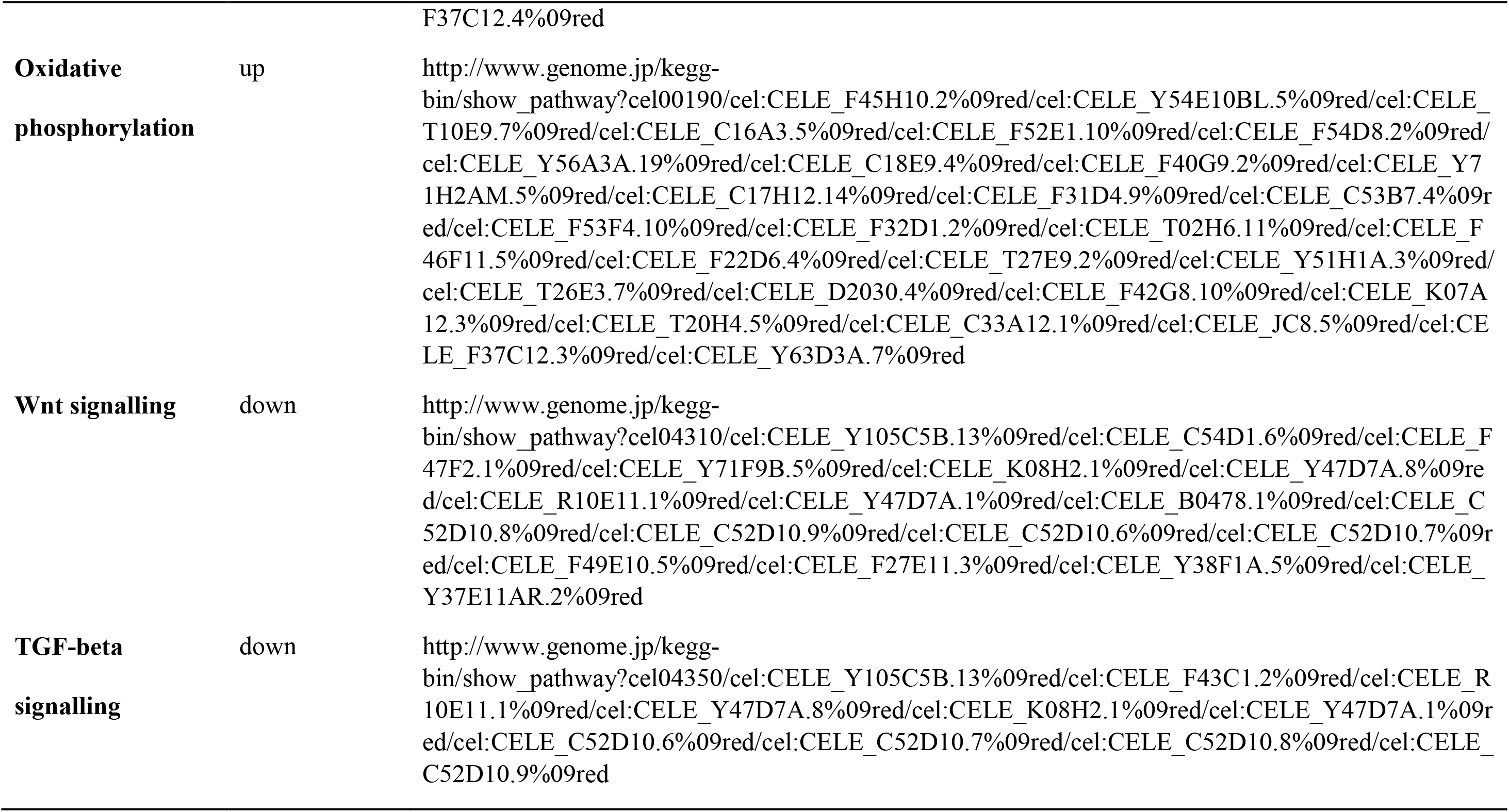
Hyperlink of significant pathway to the KEGG pathway.

**Fig. 11:**
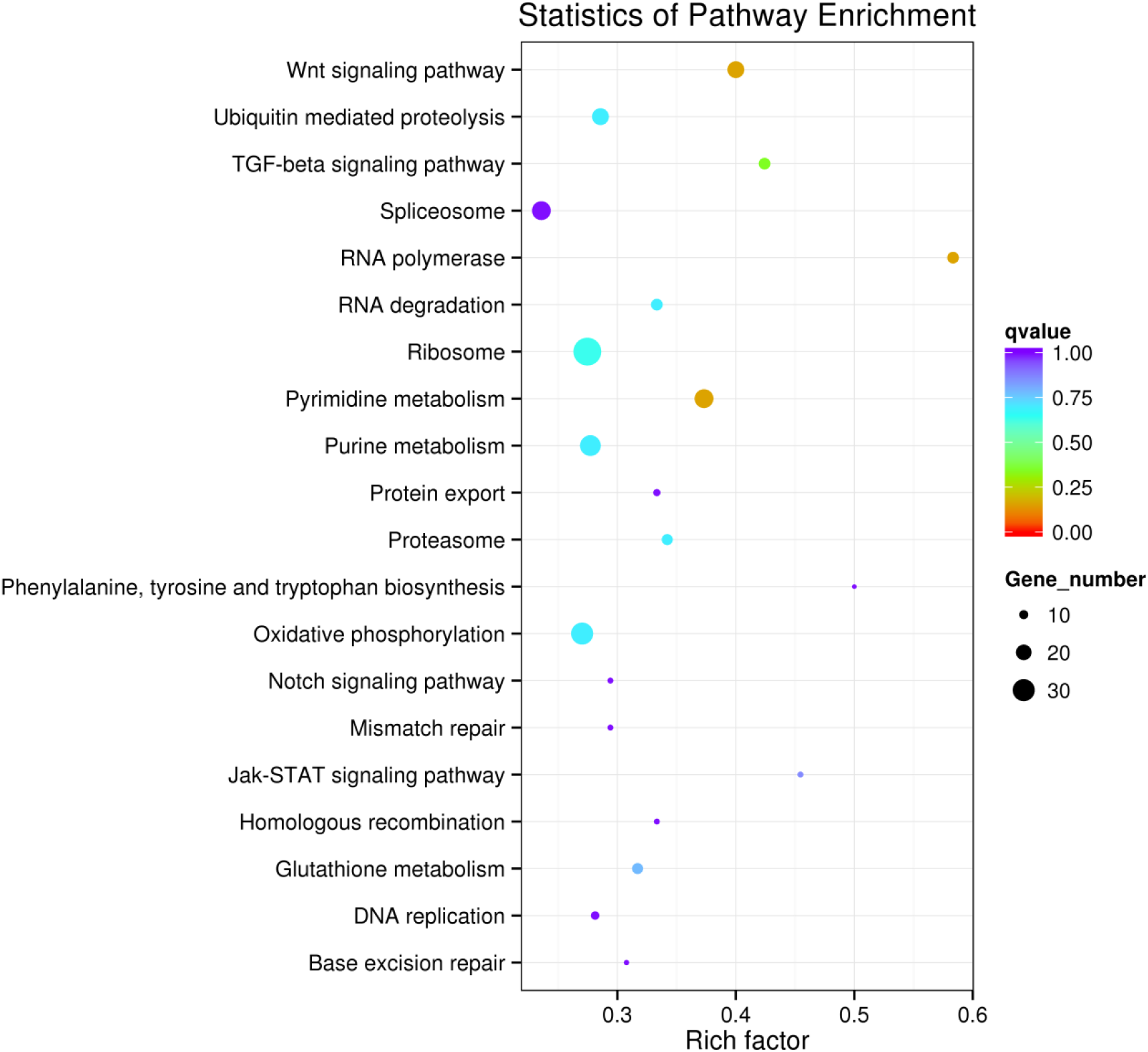
Statistics of overall KEGG pathway enrichment.

**Fig. 12:**
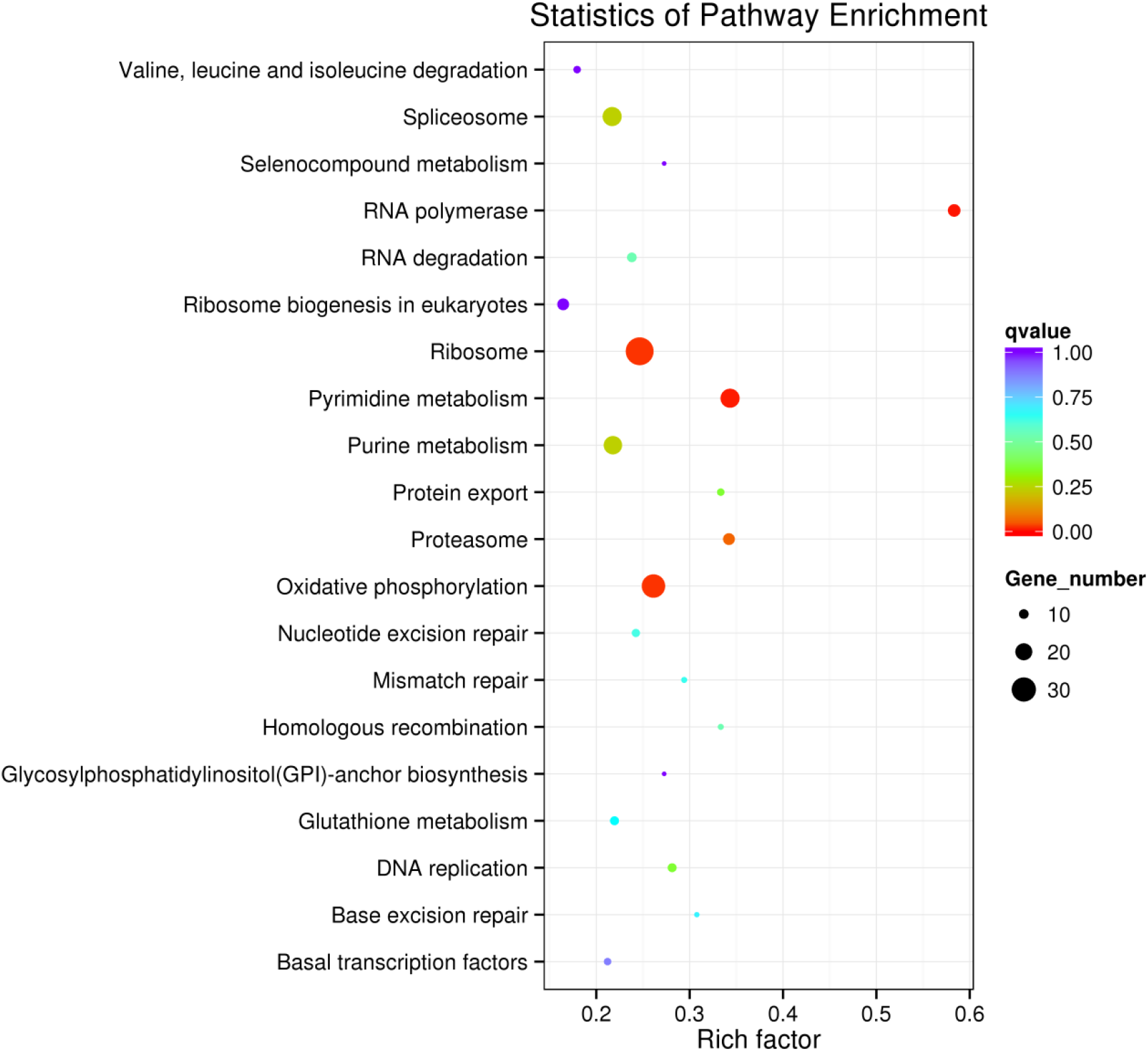
Statistics of down-regulated enriched KEGG pathway.

**Fig. 13:**
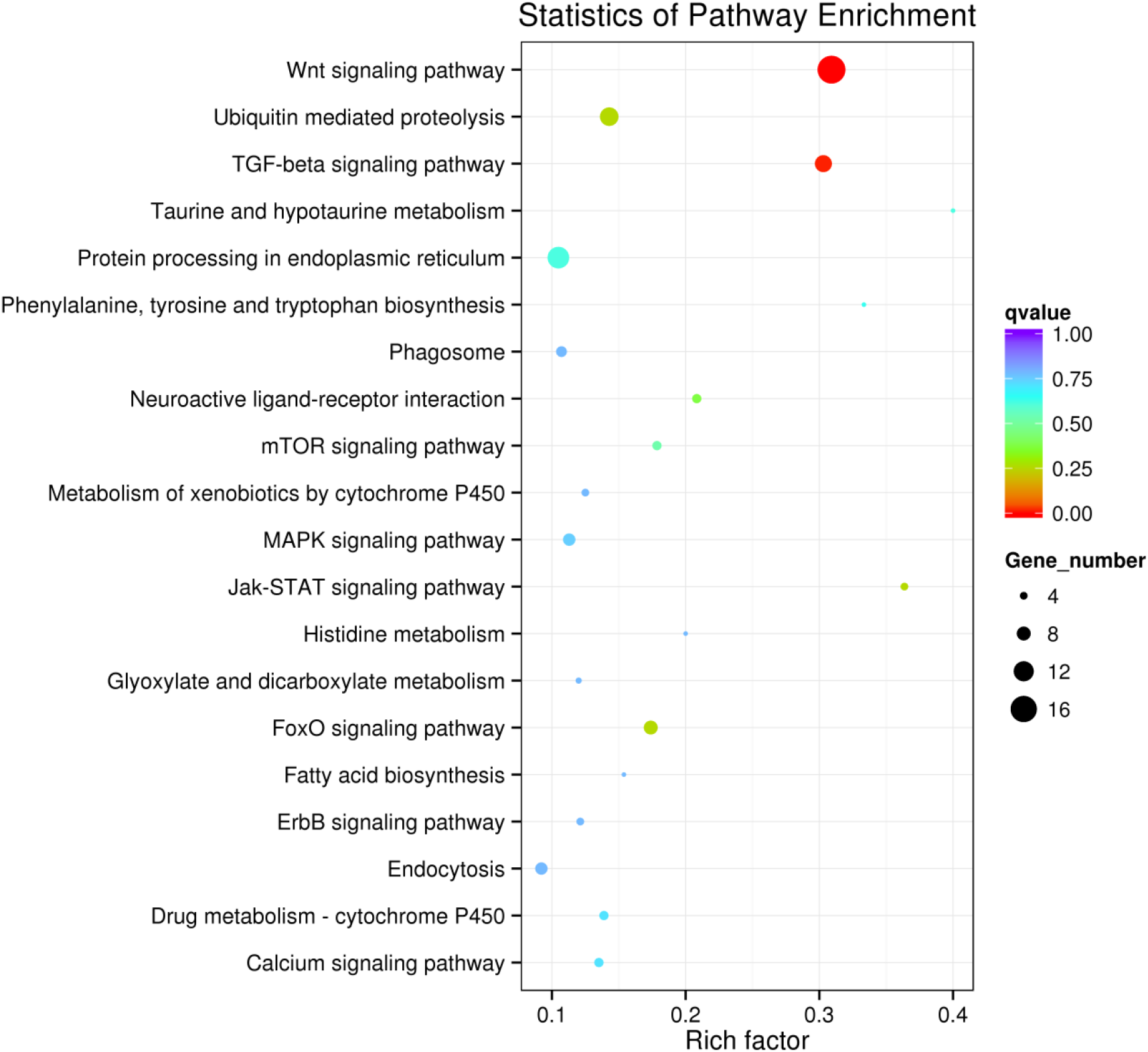
Statistics of up-regulated enriched KEGG pathway.

### Real-time PCR validation of transcriptomic results

Validation of differentially expressed genes that were identified through transcriptomic studies were performed by running qPCR analysis on 10 selected genes (5 up-regulated and 5 down-regulated genes). The result is presented in Supplementary 8.

## DISCUSSION

Patients suffering from Alzheimer’s Disease (AD) have grown in numbers with the increase in life expectancy of the world population. AD is the most frequent cause of dementia and the number of individuals afflicted with this complex disorder number is predicted to increase in the next two decades [26]. The search for effective treatments using drugs and natural products targeting multiple molecular pathways have been ongoing with little success. Disease-modifying symptom reduction therapies have encountered various shortcomings with the occurrence of adverse side effects. This drawback necessitates further screening for effective anti-AD drugs.

Danshen has been widely used for treatment of cardiovascular diseases as well as cerebral ischemia. The composition of Danshen like diterpenoid quinones and hydroxycinnamic also beneficial in treating and improved cognitive deficits in mice, protection of neuronal cells, prevention of amyloid fibril formation in AD. Sal A is one of the main active ingredients of Danshen. [27]. The structure of Sal A and curcumin are quite similar. Previous study showed that curcumin is an anti-Aβ aggregation agent [28]. Consequently, we used Sal A to observe global transcriptomic response of AD *C. elegans*.

Oxidative stress has been linked to numerous of pathologies. Antioxidant response genes forms an important part as the antioxidative defence system in combating stress posed by accumulation of ROS [29–31]. This transcriptomic study showed that three antioxidant response genes, namely *gst-4*, *gst-10* and *spr-1*, were upregulated by Sal A. These antioxidant response genes provide defence to oxidative stress caused by the production of Aβ_42._ The *sod-1* gene that was suggested to increase the life span of the *C. elegans* strain GMC101 was also upregulated.

The gene *gst-4* encodes for a stress-responsive glutathione S-transferase (GSTs) that produces resistance to oxidative stress in *C. elegans* [30]. There are 51 putative GST-encoding genes in *C. elegans* besides GST-4. GSTs are a family of enzymes that catalyse the conjugation of reduced glutathione to a wide range of substrates by deprotonating the glutathione to form thioether bonds with electrophilic atoms within the substrates [32]. The glutathione conjugational reaction serves to detoxify endobiotic and xenobiotic toxins that include products of oxidative damage [33]. Studies have shown that overexpressed *gst-4* gene led to the resistance in oxidative stress by paraquat, but does not increase life span [30]. Additionally, RNAi of *gst-4* decreases the life span of *daf-2* mutant [34].

The gene *gst-10* encodes glutathione S-transferase P 10 (EC:2.5.1.18) which has the capability to detoxify 4-hydroxynon-2-enal (HNE), a lipid peroxidation product caused by oxidative stress [29]. Overexpression of *gst-10* gene extends the lifespan of *C. elegans*. RNAi of *gst-10* increased sensitivity to HNE toxicity, and it reduced life span in both wild-type and *daf-2* mutant population [35].

The gene *spr-1* codes for a putative corepressor protein SPR-1 which acts with SPR-4, the homologue of repressor element 1-silencing transcription factor (REST) in *C. elegans* [36]. Both form part of a multiprotein complex that deacetylates and demethylates specific sites on histones. They are involved in the transcriptional repression of the presenilin protein HOP-1 [36]. Mutants of *spr-1* were shown to have reduced survival during oxidative stress [36]. This may well indicate that Upregulated *spr-1* is important for reducing oxidative stress [31]. Thus, it is suggested that Sal A not only provided neuroprotection to the *C. elegans* but also activated antioxidant response genes to diminish the oxidative stress of ROS.

According to the oxidative damage theory, it is assumed that ROS causes the accumulation of molecular damage which include superoxide (O2•−). This theory indicates that ROS which include superoxide and its derivatives causes molecular damage to accumulate and this leads to ageing [37]. The function of enzyme superoxide dismutase (SOD) is to convert O2•− into hydrogen peroxide (H_2_O_2_), which is then converted into H_2_O and O_2_ due to the presence of catalase, glutathione peroxidase, and other enzymes. According to the oxidative damage theory, SOD has the tendency to protect against ageing and also oxidative damage [37].

The effects in manipulating the expression of *sod-1*, the major cytosolic Cu/Zn-SOD isoform, suggest that cytosolic O2•− and the damage that it causes contribute to *C. elegans* ageing. Life span was shown to be slightly decreased when deletion of or RNA-mediated interference (RNAi) of *sod-1* occurs [38–41] but overexpression of *sod-1* increases life span [38]. However, it was reported that overexpression of *sod-1* did not reduce levels of oxidative damage. Furthermore, studies had shown that *sod-1* overexpression lines were hypersensitive and not resistant to oxidative stress [37]. This suggested that overexpression of SOD might not reduce ROS-induced damage in these strains.

There were reports indicating increased expression of *gst-4* led to an increase of resistance to oxidative stress, but there was no impact on lifespan [30]. Thus, this had shown that this particular antioxidant response gene is responsible in reducing the oxidative stress induced by Aβ_42_.

Based on the transcriptomic results, two genes that are involved in Alzheimer’s disease were affected by Sal A. The first one is upregulated glutathione reductase 2, *trxr-2,* gene which involved in Aβ peptide and amyloid deposits and another gene is *ptl-1* was found to be downregulated. The *ptl-1* gene is the homolog of the MAP2/MAP4/tau family in the nematode. The *trxr-2* gene was upregulated while *ptl-1* gene was downregulated.

The *trxr-2* gene has been reported to encode for a mitochondrial protein which is operated by a putative mitochondrial targeting sequence (MTS) at the N-terminus [42] and function as a thioredoxin reductase based on enzymatic assays [43]. The mitochondrial thioredoxin system under mitochondrial unfolded protein response (UPR^mito^) obviously functions to decrease incorrect disulphide bonds in misfolded proteins to ease protein refolding or exporting proteins from mitochondria to be degraded by proteasome [44]. It is essential to maintain cellular and subcellular proteostasis so that the organism can survive. Its imbalance may jeopardize the function of all cellular organelles which include mitochondria and this will results in enhanced progression of ageing and development of Alzheimer’s disease [45].

The *trxr-2* gene was known to have a protective role on transgenic *C. elegans* CL2006 strain. It was reported that downregulation of *trxr-2* resulted in enhanced paralysis and TRXR-2 protein expression when increased, did not attenuate the paralysis onset [42] Overexpression of TRXR-2 prominently lessened total levels of Aβ species and amyloid deposits by degrading most of Aβ load. This might be due to the interaction of TRXR-2 with proteins that are involved in Aβ peptide and amyloid deposits degradation in the mammals. The proteins responsible in reducing Aβ peptide and amyloid deposits could be angiotensin-converting enzyme, insulin-degrading enzyme or neprilysin. These orthologs of these enzymes are found in the nematodes [42]. Another possibility is that overexpression of TRXR-2 causes insulin signalling to diminish, which is then led to Aβ autophagic-dependent degradation [42]. The molecular mechanisms of TRXR-2 in attenuating paralysis and reducing amyloid deposits in Aβ nematodes, may pose as a promising application of knowledge to the human scenario suffering in AD [42].

Another gene affected by Sal A was downregulated of *ptl-1* gene. This gene encodes protein with tau-like repeats. The PTL-1 protein was found to acquire strong homology in sequence to tau/MAP2/MAP4 family of MAPs over the repeat region [46, 47]. Furthermore, PTL-1 shares similarities with tau protein in other aspects like size, amino acid content, charge distribution, predicted secondary structure, hydrophobicity and flexibility. These two proteins also have numerous potential glycosylation sites and many phosphorylation sites [47]. In humans, tau [neuronal microtubule associated protein (MAP)] hyperphosphorylation leads to abnormal self-assembly of tangles of paired helical filaments and straight filaments, which is associated to Alzheimer’s disease [48]. The *ptl-1* gene functions to perpetuate normal neuronal health through cell-autonomous mechanism and regulate animal lifespan [49]. The functions of this gene also include the involvement in cholinergic/GABAergic transmission, mechanosenstation and microtubule assembly lifespan [49]. With the effect of Sal A in reducing the expression of *ptl-1* gene, it may work well in the human scenario in decreasing the mutated tau expression that causes Alzheimer’s disease.

Besides upregulating antioxidant response genes, there were ten genes that were identified to be top upregulated differential expressed genes. These genes include genes that encode for FIP (Fungus-Induced Protein) Related (NP_741606), major sperm protein (NP_499807), C-type Lectin (NP_503650) and Serpentine Receptor, class T (NP_504173). The rest of the genes were hypothetical genes.

The gene *T05D4.5* encodes a major sperm protein which plays a significant role in the motility machinery of *C. elegans* by driving the crawling movement of the mature sperm. Furthermore, it functions as a hormone on the oocytes of the female worms by triggering the contraction of the oviduct wall to allow oocytes to be fertilized and thus allowing their maturation [50]. As for the C-type lectin, it is a carbohydrate-binding protein that has various functions such as cell adhesion, immune response to microorganisms and sensing apoptosis [51–53]. Serpentine receptor which is encoded by *srt-13* gene is also known as G-protein-linked receptor. The receptor acts as a mediator in responses to various signal molecules like hormones and neurotransmitters [54]. The gene *fipr-11* which encodes for FIP (Fungus-Induced Protein) is believed to encode for possible antimicrobial peptides (AMPs) [55]. It is interesting to note that the antimicrobial activities of AMPs act as a defense system for *C. elegans*. AMPs work by disrupting the anionic cell walls and phospholipids membranes of the microorganisms [55]. Taken together, Sal A might have the potential in upregulating the transcription of genes that were involved in reproduction, immune response to microbes and antimicrobial activities.

Among the top downregulated genes that were affected due to the presence of Sal A were *acr-23*, *daf-7* and *cpr-2* which encoded for betaine receptor ARC-23, dauer larva development regulatory growth factor DAF-7 and cysteine protease related protein, respectively. The betaine receptor ACR-23 functions as a ligand-gated cation channel in *C. elegans*. The protein ACR-23, which is active when betaine is present, plays a role in mechanosensory neurons that regulate locomotion of the nematode [56]. It was reported that *acr-23* was highly expressed in six mechanosensory neurons, multiple interneurons and body muscles. The ACR-23 protein was highly expressed in muscle in larvae but lowly expressed in adults. Furthermore, *C. elegans* mutant strains that have deleted *acr-23* gene demonstrated mild swimming defects, crawling sluggishly on the agar plate and crawling was sometimes disrupted with repeated pauses [56]. The gene *daf-7* that was annotated for dauer larva development regulatory growth factor TGF-β. DAF-7 expression levels are regulated due to the presence of food but downregulated by pheromones. DAF-7 functions through a heteromeric TGF-β receptor that consisted of DAF-1 and DAF-4 that have impact towards the activity of the transcription factors DAF-8 and DAF-14 [57]. DAF-7 is believed to modulate the activity of neural circuits in a duration period of time. Another possible function of DAF-7 is that it acts as a neuromodulator to some chemosensory neurons that works synergistically or antagonistically [58]. As for *cpr-2* gene, it encodes for cysteine protease that breaks down protein by attacking the peptide-bond after cysteine residue which acts as a nucleophile is activated by histidine residue [59].

The other downregulated proteins that include BBS-1, CEH-23, CLH-4, CFZ-2, CYD-1, SUE-1 and FBXB-17 were annotated as Bardet-Biedl syndrome 1 protein homolog, Homeobox protein CEH-23, Chloride channel protein, *Caenorhabditis* FriZzled homolog, Suppressor of Elongation defect protein and G1/S-specific cyclin-D respectively. The gene *bbs-1* in humans is responsible for Bardet-Biedl syndrome, an obesity syndrome with symptoms of obesity, mental disorder, hypogenitalism, retinal degeneration, renal failure and short stature [60]. Since *bbs-1* in *C. elegans* is a homolog to the human gene *bbs-1*, the repression of the *bbs-1* transcription indicated that Sal A may disrupt the function of the gene in humans. As for *clh-4* gene, it is one of the CLC-type chloride channel genes in the nematode. As opposed to human who has nine different identified CLC genes, the worm has only six identified genes. The probable roles of CLC Cl^-^ channels are to stabilize membrane potential, undergo transepithelial transport, regulate cell volume, and conduct endocytosis [61]. The protein expression of CLH-4 normally occurs in the large H-shaped excretory cell [62, 63]. The excretory cell has been shown to maintain osmotic balance and internal hydrostatic pressure in the worm [64]. Worms without an excretory cell bloat could not survive within 24 hours. Moreover, this particular cell is sensitive to changes in external osmolarity [64]. As for *cfz-2* gene, it plays a role in cell migration, axon development, organization of the anterior ganglion [65]. Furthermore, it acts as a receptor for Wnt proteins [66, 67].

In the nematode, the *cyd-1* gene which cooperates with *cdk-*4 gene, regulates the postembryonic developmental stage of G1 phase of the cell cycle [68–70]. Working together with *cdk-4* gene, *cyd-1* also participates in sex determination during gonadogenesis. This is done by influencing the somatic gonadal precursor cell to divide asymmetrically [71]. Moreover, the *cyd-1* gene uniquely affects the growth of the coelomocyte lineage and intestinal cells during late embryogenesis [68, 72]. The specific functions of *sue-1* and *fbxb-17* which encode for SUpressor of Elongation defect and F-box B proteins respectively, are currently unknown. Taken together, Sal A downregulated the gene transcription that were responsible for locomotion, ligand-gated cation channel and neuromodulation of chemosensory neurons.

Gene Ontology (GO) term enrichment is an analysis in which genes are assigned to a set of hierarchical classifications depending on their functional characteristics. It is performed to categorize differentially expressed genes (DEGs) into according to their biological functions [73]. In this study, GO analysis showed that Sal A upregulated mostly genes that were involved in cellular component as compared to the downregulated genes. When compared to upregulated genes, Sal A downregulated most of the genes that were responsible for biological process and molecular function.

As for the KEGG analysis, Sal A significantly upregulated genes that were involved in RNA polymerase, pyrimidine metabolism, ribosome and oxidative phosphorylation pathways. These pathways play important roles in gene regulation. In RNA polymerase pathway, genes that were associated with RNA polymerase I, II and III were upregulated.

Pathways such as ribosome and oxidative phosphorylation are associated with energy supply and metabolism of materials. Oxidative phosphorylation involves the usage of enzymes in the cells to oxidize food for energy to produce adenosine triphosphate (ATP). The site where oxidative phosphorylation occurs is electron transport chain which is in the mitochondrion of the *C. elegans* [59]. The electron transport chain involves five complexes that are important for respiration of the nematode. These include NADH-coenzyme Q oxidoreductase (complex I), Succinate-Q oxidoreductase (complex II), Electron transfer flavoprotein-Q oxidoreductase, Q-cytochrome c oxidoreductase (complex III) and Cytochrome c oxidase (complex IV) [74]. In this study, Sal A triggered the genes that involved in NADH-coenzyme Q oxidoreductase (complex I), cytochrome c oxidoreductase (complex III) and Cytochrome c oxidase (complex IV) to be upregulated. The NADH-coenzyme Q oxidoreductase complex involves the removal of electrons from NADH and transferred to CoQ [74]. As for cytochrome c oxidoreductase complex, CoQH_2_ is reduced due to oxidized NADH or succinate that cause the transfer of two electrons to produce oxidized CoQ. In cytochrome c oxidase complex, the reduced cytochrome *c* is due to transfer of electrons to copper ions (Cu_a_^2+^), subsequently to cytochrome *a*, then to another copper ion (Cu_b_^2+^) and cytochrome *a_3_* and last but not least, to O_2_ to produce H_2_O [74].

Ribosome, which has both large and small rRNAs, acts as a site for protein synthesis to occur [74]. In this study, Sal A triggered the upregulation of genes in RNA polymerase, pyrimidine metabolism, oxidative phosphorylation and ribosome pathways. Hence, this might suggest that the production of antioxidant response proteins was increased to act as defensive system towards ROS produced by Aβ.

Genes were downregulated due to Sal A were involved in Wnt-signaling pathway and TGF-beta signaling pathway. The Wnt signalling pathway is consisted of signal transduction pathways that involves the passing of protein signals into a cell *via* cell surface receptors. There are three types of Wnt signalling pathway which include canonical Wnt pathway, the noncanonical planar cell polarity (PCP) pathway, and the noncanonical Wnt/calcium pathway. The canonical Wnt pathway, controls the transcription of the specific target genes with the aid of β-catenin protein [75]. In noncanonical planar cell polarity pathway, the cytoskeleton which plays significant role of cell shape is regulated. The noncanonical Wnt/calcium pathway is responsible in regulating calcium in the cell [76, 77].

Wnt signaling is responsible in regulating various cell processes throughout the embryonic development of *C. elegans*. Wnt signals are important in regulating cell fate specification, cell division and cell migration [78]. As for the transforming growth factor beta (TGFB) signaling pathway, it plays important role in the development of the postembryonic mesoderm which include antero-posterior pattern of the nematode [79]. With genes shown to be downregulated in Wnt signaling and TGFB signaling pathways, it can be assumed that Sal A influences both the embryonic and postembryonic developmental stages of the worm. With regards to the results obtained, Sal A may act as a disease modifying drug for Alzheimer’s disease.

## Conclusion

In this study, we found that Sal A treatment towards *C. elegans* expressing human Aβ_42_ gene showed positive response. From transcriptomic analysis, the antioxidant genes in *C. elegans* were up-regulated when treated with Sal A. In addition, the significantly expressed Alzheimer’s disease related genes were all respond accordingly to rescue the *C. elegans* from Aβ_42_ toxicity. Through this study, we infer that potential Sal A has the potential to be an alternative drug to combat Alzheimer’s disease.

## Acknowledgements

We would like to thank all our collaborators and colleagues for the discussion and the work conducted in this lab. This study was funded by the USM Top Down Research Fund – URICAS (1001/PBIOLOGI/870029).

